# A novel homecage operant paradigm reveals circadian and behavioral dynamics of social motivation in mice

**DOI:** 10.1101/2025.08.21.671364

**Authors:** Yiru Chen, Susan E. Maloney, Alexxai V. Kravitz

## Abstract

Deficits in social motivation are a core feature of many neurodevelopmental disorders, including autism spectrum disorders. While a range of tools have been developed to quantify social motivation in rodents, most rely on brief tests in dedicated apparatuses that can introduce stress and novelty, potentially reducing test reliability. Most current approaches are also typically not suited to studying learning across days or circadian rhythms of social motivation. To address these challenges, we developed a social operant task that can run around-the-clock for multiple days in the mouse homecage, continuously monitoring social motivation around the circadian cycle. The task consists of a custom-built automated door that was installed between two rodent homecages and configured so one mouse could trigger the door to open with a nose-poke action from within their cage. When open, the door allows for social interaction with the neighboring mouse through a perforated stainless-steel panel, which did not allow the mice to cross over into the neighboring cage. Mice opened the door multiple times each day, allowing us to quantify their amount and daily rhythms of social motivation. In our first experiment, C57BL/6J mice (both sexes) were individually housed with an empty adjacent cage for five days, after which a same-sex social partner was introduced for another nine days. Mice opened the door at significantly higher rates when the social partner was present vs. absent, confirming that mice were motivated to earn social interaction. This task also revealed a circadian rhythm to social motivation that peaked about 2 hours after the peak in their feeding rhythm. We speculate that mice first addressed their caloric needs each day before changing their priority to social behavior. Given prior literature implicating the dopamine system in social motivation, we also tested whether dopamine antagonists would block social motivation in our task in a new group of 14 mice (both sexes). The dopamine D1 receptor antagonist SCH23390 (delivered systemically at 0.3mg/kg SC) reduced social seeking without affecting locomotor activity or food intake, demonstrating a selective role for dopamine D1 receptors in social motivation. The dopamine D2 receptor antagonist haloperidol (delivered systemically at 0.3mg/kg SC) also reduced social seeking, but reduced locomotor activity and food intake as well, demonstrating a general reduction in behavior that was not specific to social motivation. Overall, our task offers a way for studying social motivation in the rodent homecage, which has advantages for studying disorders that involve both social and circadian disruptions.

## Introduction

Social motivation, the drive to initiate and maintain social contact, is a fundamental component of behavior across species and is essential for survival and mental well-being^1–3^. In humans, disruptions in social motivation are suggested to underlie the social challenges that hallmark several neurodevelopmental and psychiatric disorders, including autism spectrum disorder (ASD)^3^, schizophrenia^4^, and depression^5,6^.

Behavioral tests in rodents have uncovered many neural and molecular underpinnings of social motivation. These behavioral tests can be broadly broken into three main categories: social interaction, social approach, and social operant tasks. Social interaction tests include the dyadic interaction test, where two mice are allowed to explore one another, and social behaviors such as sniffing or allogrooming are quantified^7^. Social approach tests include the 3-chambered test, in which mice are given the option to investigate a novel mouse or a novel object^8,9^. This test can measure the relative time investigating and the number of approaches to the mouse vs the object, which is indicative of sociability and social preference. Despite the power of social interaction and approach tests, neither class is suited to quantifying how motivated a mouse is for social interaction, cost-benefit decisions about social drive vs. other drives, or learning across days. Social operant tasks were developed to address these questions about social motivation. In a social operant test, an animal is trained to perform an action (typically a lever-press or a nose-poke response) to gain access to a conspecific^8–11^. The number of times a mouse performs the action, or “works”, to obtain social access is interpreted as a proxy for their level of social motivation.

Social operant paradigms have provided key insights into the neural and molecular basis of social motivation. For example, Hu et al. (2021) identified an amygdala-to-hypothalamus circuit involving projections from the basomedial amygdala to the medial preoptic area that promotes social motivational behavior. This pathway was shown to influence the activity of ventral tegmental area (VTA) dopamine neurons, providing a link between limbic social processing and dopaminergic motivation circuits^9^. Ramsey et al. (2023) showed that female CD-1 mice exhibit significantly stronger social operant self-administration and a preference for social interaction over palatable food compared to female C57BL/6J mice, highlighting strain-dependent variation in social motivation^10^. Our recent work revealed sex-specific deficits in social motivation in *Shank3b* mutant mice, and that blocking oxytocin receptors reduced social reward seeking in C57BL/6J mice^11^, further demonstrating the power of the social operant paradigm to dissect both circuit-level and genetic contributions to social behavior.

However, social operant tasks typically utilize specialized chambers, into which mice are transported for daily testing. Moving mice into new environments can introduce both stress and novelty, which can interfere with behavior^12,13^ and confound interpretations of the task. For instance, it is not always clear whether operant responses truly reflect a desire for social interaction or reflect test chamber discomfort or exploration of the new environment. In addition, these paradigms often require several days of training before data collection can begin, and testing sessions are typically limited to short durations (e.g., one hour per day), which restricts data yield and may miss critical temporal dynamics, such as circadian patterns of social motivation.

To address these limitations, we developed a novel open-source home-cage-based social operant paradigm to assess social motivation without removing mice from their home-cages. Our system is automated, which allows us to assess social motivation for several consecutive days without experimenter intervention, reducing stress and increasing the scale of data collection. The system consists of a battery-powered servo-driven social door that provides access to mice in the neighboring cage, which is controlled by a Feeding Experimentation Device version 3 (FED3)^14^. The door also integrates proximity sensors that report whether Test and Stimulus mice approach the door after each social operant response, allowing us to distinguish social seeking from general exploration of the FED3 device. As our paradigm runs for multiple days, we can assess social motivation across the circadian cycle to determine when social motivation peaks and how it relates to other daily rhythms of behavior. Finally, we have released all design files online, along with instructions on how to build the system. Taken together, our paradigm offers a powerful and flexible platform for investigating social motivation in rodent home cages.

## Methods

### Animals

C57BL/6J (JAX: 000664) male and female mice were bred in-house. Upon weaning at postnatal day (P)21, mice were group-housed according to sex. All mice were housed on a 12-hour light/dark cycle (lights on at 6:00 AM) with room temperature (∼20-22°C) and relative humidity (∼50%). During testing, both test and stimulus animals were single-housed in ventilated (28.5 × 17.5 × 12 cm) translucent plastic cages with corncob bedding, nestlet squares, food pellets (Dustless Precision Pellets®, Rodent, Grain-Based, 20 mg; Bio-Serv, Product #F0163), and water provided ad libitum. The bedding was changed when necessary, with a maximum time of 14 days between cage changes. Weights were recorded daily between 2-3 pm. All behavioral tasks were conducted by a female experimenter. All procedures were approved by and performed in accordance with the Institutional Animal Care and Use Committee at Washington University in St. Louis and conformed to NIH guidelines for the care and use of laboratory animals.

In the validation experiment, we used two independent cohorts of mice, which were run serially to accommodate equipment availability. The first cohort contained 9 test mice (4 males and 5 females, postnatal day 50–55) and 9 same-sex stimulus mice (P53–56 when introduced) in the first cohort. The second cohort contained 6 test mice (3 males and 3 females, P50–55) and 6 same-sex stimulus mice (P48–60 when introduced) in the second cohort.

In the dopamine antagonist experiment, we also used two independent cohorts of mice. The first cohort contained 7 test mice (4 males and 3 females, 11-14 weeks old) and 7 same-sex stimulus mice (P85 when introduced) in the first cohort. The second cohort contained 10 test mice (5 males and 5 females, mostly 15-16 weeks old, one female was 20 weeks old) and 10 same-sex stimulus mice (mostly 18-20 weeks old, one female was 22 weeks old) in the second cohort. Before Experiment 2, 8 mice (4 males and 4 females, 14 weeks old) were used in the pilot test to determine the appropriate dosage of dopamine receptor antagonists.

### Homecage Social Operant Assay

The social home-cage operant paradigm consisted of three components: 1) a pair of home cages (28.5 × 17.5 × 12 cm), 2) a custom social door, and 3) a FED3 device. A 5.4cm hole was drilled into each home-cage, and the social door was installed between the cages to control social interactions between the cages (Figure 1A–D, Supplemental Figure 1). FED3 is a smart pellet dispensing device that includes two nosepoke ports for mice to interact with and a pellet dispenser to quantify patterns of food intake^14^. FED3 was programmed to send a 500- ms 3V pulse to the social door each time the Test mouse poked on the active port. This pulse triggered the social door to open for 12s (Figure 1D). The other port was inactive, and a poke within did not result in an outcome. The mouse that controlled the door opening via FED3 was termed the “Test” mouse, and the mouse in the target cage was termed the “Stimulus” mouse. To minimize the risk of injury, the door also made several small movements to warn the mice before it closed. Once open, the Test and Stimulus mice could interact through a stainless-steel grate that was perforated with five 1-cm holes, which enabled visual contact and limited physical interactions (e.g., nose touching and sniffing), but did not allow the mice to cross between the cages. Two Time-of-Flight (ToF) proximity sensors were used to track the behavior of the Test and Stimulus mice while the door was open (Figure 1D). Nosepokes, feeding events, door openings, and proximity data were stored on SD cards and retrieved after the experiment.

**Figure 1.**
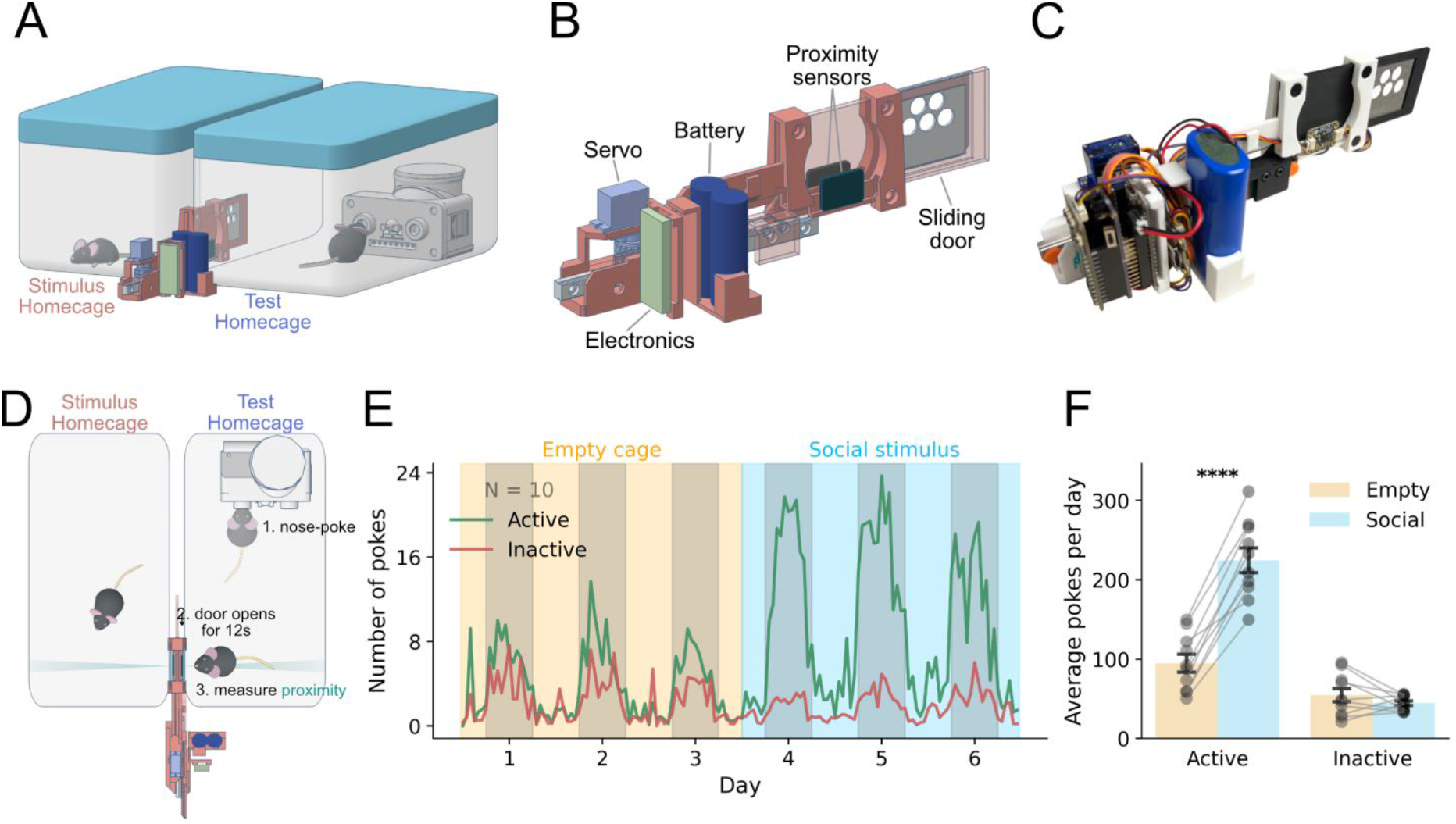
A social homecage operant paradigm for assessing social motivation in a low-stress environment. (A) Setup showing the operant device integrated into adjacent homecages, allowing voluntary social interaction through an automated door. (B) Schematic of the operant device featuring a nosepoke port, sliding door, proximity sensors, and circuitry for automated control. (C) Photograph of the physical device including servo motor, battery, and sensor components. (D) Behavioral sequence: nosepoke triggers door opening, granting access to an adjacent chamber for social approach measurement. (E) Behavioral data showing operant nosepoke responses across 6 days for 10 mice, divided into an empty cage period and a same-sex social stimulus period. Active and inactive nosepokes are plotted separately. Gray shading represents the dark phase. Mice showed a sharp increase in active pokes following the introduction of a social partner. (F) Quantification of average daily active and inactive pokes reveals a significant increase in active responses for access to a social partner (N = 10, p = 1.08 × 10^−6^, paired t-test), with no change in inactive pokes (p=0.23). Asterisks indicate statistical significance: p < 0.0001 (****). All tests were two-tailed, and multiple comparisons were corrected using the Bonferroni method where applicable.

### Assembly of the social door

The social door system consisted of a 3D-printed frame and sliding door, a servo motor (Feetech FS90R), a PCF8523 real-time clock (RTC, Adafruit 5189), a limit switch (HiLetgo 3-01-1546), two VL6180X ToF proximity sensors (Adafruit 3316), and a 4400 mAh rechargeable battery (Adafruit 356). Arrangement of components is shown in Figure 1A and Supplemental Figure 1A. A custom printed circuit board (PCB) connected these components to an Adafruit Feather M0 Adalogger (Adafruit 2796), which controlled door movement and interfaced with the sensors and an OLED screen module (Adafruit 4650) to display real-time information such as battery level, proximity values, and system status (Supplemental Figure 1B–C). A portion of the sliding panel was covered with a laser-cut perforated stainless steel plate through which mice could interact (five centered holes with a diameter of 1 cm, Figure 1A–B). A 5.4 cm-diameter hole was drilled into the side of the test and stimulus cages to provide an interaction port for the door. Magnets were used to connect and align the door to the two cages.

### Quantification of circadian rhythms of feeding

Feeding was assessed concurrently with social behavior using the FED3 device^14^. Pellets were dispensed on a “free feeding” schedule, such that the pellet (20mg grain-based pellet, BioServ) was replaced each time it was removed, and FED3 logged times of each pellet retrieval on its internal SD card for later analysis.

### Quantification of circadian rhythms of locomotion

Locomotor activity was continuously monitored using battery-powered in-cage motion sensors (Pallidus MR1). These sensors use a passive infrared (PIR) sensor to detect locomotion in the cage and upload it to a cloud database using LoRaWAN. Data was viewed on the cloud, and raw data was downloaded for analysis.

### Administration of Dopamine Receptor Antagonists

Mice were single-housed and linked to an empty cage for four days (Figure 5A). During this time, they received saline injections on Days 2 and 4 to acclimate them to the injection procedure. A social partner was introduced on Day 5, followed by randomized administration of saline or dopaminergic antagonists every 2 days. D1 receptor antagonist SCH23390 (0.3mg/kg SC) and the D2 receptor antagonist haloperidol (0.3mg/kg SC) were injected 1.5 hours before dark onset, and behavior was quantified during the first four hours of the dark cycle. Dosages were determined in pilot tests using single-housed mice monitored with motion sensors, optimizing doses that caused minimal reductions in locomotion (Supplemental Figure 2). On Days 6, 8, and 10, mice received randomized injections of saline, SCH23390, or haloperidol. Treatment order was counterbalanced: 5 mice received saline → SCH23390 → haloperidol; 5 mice received SCH23390 → haloperidol → saline; and 4 mice received haloperidol → saline → SCH23390.

### Data Analysis and Statistics

Sample sizes were determined based on pilot studies and subsequent power analysis. Mice were randomly assigned to treatment groups with randomized drug injection orders. Behavioral analysis and figure plotting were conducted with Python 3.13 using Visual Studio Code. Data are presented as histograms, polar plots, box plots, or mean ± SEM plots. In box plots, the center line indicates the median, and the box edges represent the 25th and 75th percentiles. Circadian rhythms of behavior (seeking, exploring, feeding, activity) were analyzed using cosine fitting to extract acrophase^15^. Jupyter notebooks containing analysis code are provided at https://github.com/KravitzLab/Chen2025.

Data distributions were examined for the fit of parametric assumptions. Statistical tests included repeated measures ANOVAs, two-sided paired t-tests, post hoc pairwise t-tests, Friedman tests, and Wilcoxon signed-rank tests, as appropriate. Holm-Bonferroni correction was applied for multiple comparisons, where appropriate. Asterisks on the figures indicate statistical significance: p < 0.05 (*), p < 0.01 (**), p < 0.001 (***), and p < 0.0001 (****).

### Data exclusions

Overall, 32 mice were tested for social motivation in these studies, and data from 8 (25%) were excluded from analysis. These included data from 5 mice (3 males and 2 females) in our validation experiments and from 3 mice (2 males and 1 female) in the dopamine antagonist experiments. The reasons for exclusion included device failures of the social door or failure of the mouse to use the FED3 device to obtain food. In addition, data from specific days of a few animals were excluded due to temporary device failures.

## Results

### The homecage social operant test

Ten Test mice (4 males and 6 females) underwent three experimental phases: (1) an empty-cage period (5 days) during which valid pokes opened the door to an adjacent empty cage, (2) social stimulus period (9 days) during which a same-sex stranger mouse was introduced into the adjacent cage and replaced every 3 days with a new same-sex mouse to maintain novelty, (3) extinction period (4 days) during which the social partner was removed so the door opened to an empty adjacent cage. When comparing the last three empty-cage days and first three social days, two-way repeated measures ANOVA revealed a significant main effect of period (F(1, 9) = 59.4, p = 2.97 × 10^-5^, η^2^_p_ = 0.47), a significant main effect of poke type (F(1, 9) = 79.0, p = 9.44 × 10^-6^, η^2^_p_ = 0.75), and a significant interaction between period and poke type (F(1, 9) = 142.5, p = 8.06 × 10^-7^, η^2^_p_ = 0.55). Test mice significantly increased their active nose-poking behavior during the first 3 days of the social stimulus period, compared to when the prior 3 days, when the adjacent cage was empty (t(9) = 11.53, p = 1.08 × 10^-6^, Cohen’s d = 3.65, paired t test) (Figure 1E–F). This demonstrates both task acquisition and the ability to quantify social motivation.

We analyzed how feeding behavior varied across the Empty, Social, and Extinction periods of the operant task (Supplemental Figure 3) using a one-way repeated-measures ANOVA. This revealed a significant main effect of period on pellet retrieval (F(2, 18) = 5.23, p = 0.016, η^2^ = 0.17). Post hoc pairwise comparisons (Holm-Bonferroni correction) indicated that the number of pellets retrieved during the Extinction phase was significantly lower than during the Empty cage period (t(9) = 3.10, p = 0.038, Cohen’s d = 0.98), reflecting a reduction in reward-seeking. In contrast, there were no significant differences between the Empty and Social phases (p = 0.24), nor between Social and Extinction (p = 0.10).

### Identification of social seeking vs. exploration trials

Two Time-of-Flight (ToF) proximity sensors were used to measure the latency from the Test mouse nose-poke to both the Test and Stimulus mice approaching the door (Figure 2A–B). Test mice approached the door within 4s of their nose-poke on nearly all trials (85.7%), which we defined as “Seeking” trials (Figure 2A–B). Trials where the Test mouse did not approach the door within 4s were defined as “Exploring” trials (Figure 2A–B). A two-way repeated measures ANOVA revealed a significant main effect of period (F(2, 18) = 49.3, p = 1.13 × 10^-7^, η^2^_p_ = 0.57), a significant main effect of trial type (F(1, 9) = 200.3, p = 1.87 × 10^-7^, η^2^_p_ = 0.78), and a significant interaction between period and trial type (F(2, 18) = 61.1, p = 4.07 × 10^-7^, η^2^_p_ = 0.59). Test mice exhibited a significant increase in Seeking trials when Stimulus mice were present (t(9) = 11.7, p = 2.85 × 10^-6^, Hedges’ g = 4.17, pairwise post hoc test with Holm-Bonferroni correction), with a gradual decline across the extinction phase (t(9) = -8.99, p = 1.73 × 10^-5^, Hedges’ g = 2.23) (Figure 2C–D). In contrast, the number of Exploring trials was not impacted by the presence of the Stimulus mouse (Figure 2C–D, Supplemental Figure 4A–B).

**Figure 2.**
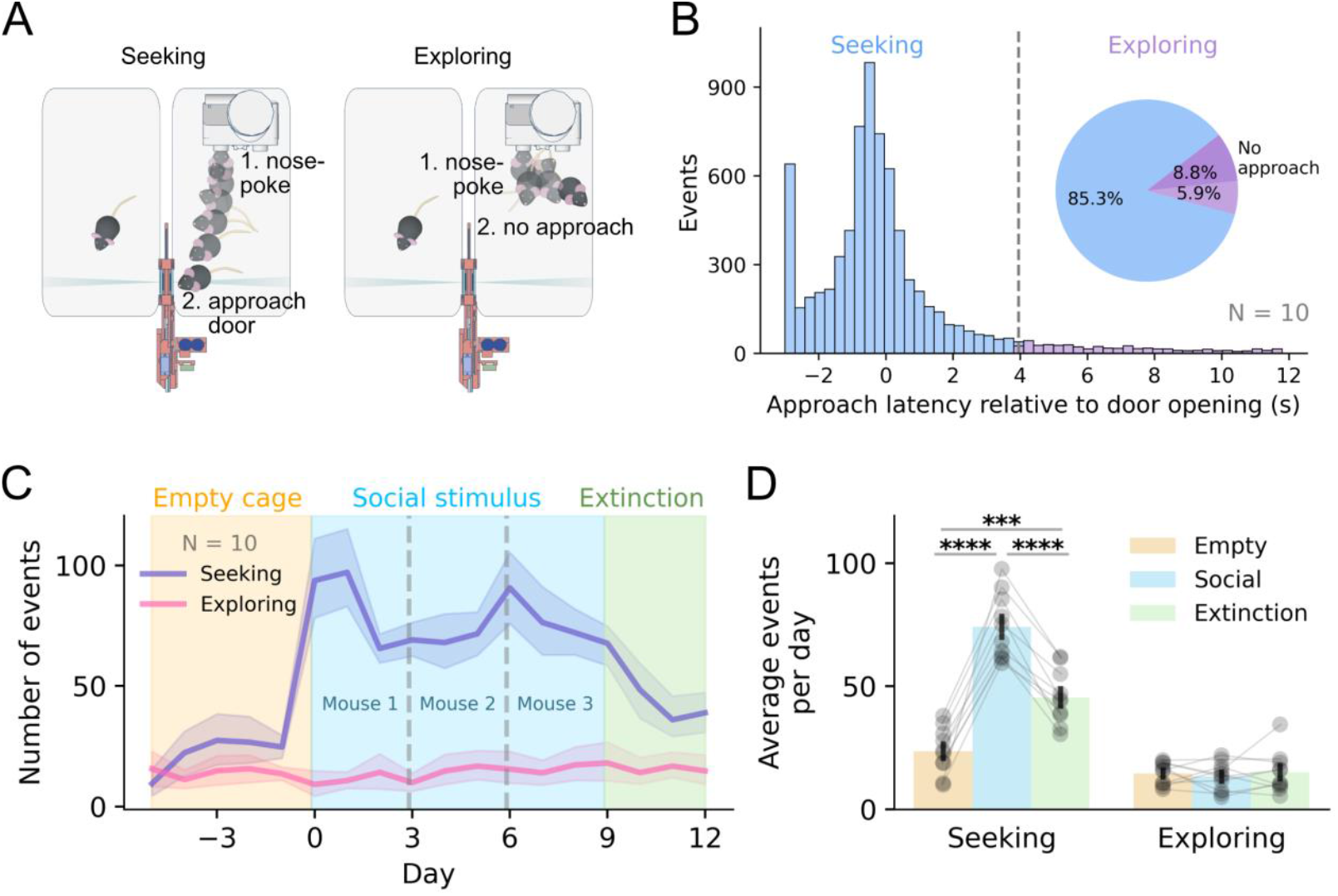
Operant social behavior reveals distinct seeking and exploring responses following door opening. (A) Schematic illustrating two post-door-opening behavioral responses: seeking, defined as approach to the door after it opens, and exploring, defined as non-approaching behavior following door fully opening. (B) Histogram of approach latency relative to door opening during social stimulus period, compiled across all trials (N = 10 mice). The majority of approaches occurred within the first 4 seconds after the door opened and were classified as seeking events (blue). Trials where mice did not approach within 4 seconds were classified as exploring (purple). The pie chart summarizes the proportions of each outcome: seeking (85.3%), delayed approach (5.9%), and no approach (8.8%). (C) Temporal profile of seeking and exploring events across experimental phases: baseline with empty cage, social stimulus exposure (3 novel same-sex mice), and extinction (empty cage of the last stimulus mouse). (D) Quantification of average events per day shows a significant increase in seeking during social stimulus days compared to both empty cage (N = 10, p = 2.85 × 10^-6^, pairwise post hoc test with Holm-Bonferroni correction) and extinction (p = 1.73 × 10^-5^), and a significant difference between empty cage and extinction (p = 9.83 × 10^-4^). Exploring behavior showed no significant change across conditions. Asterisks indicate statistical significance: p < 0.001 (***) and p < 0.0001 (****). All tests were two-tailed, and multiple comparisons were corrected using the Bonferroni method where applicable.

A social interaction was defined as the presence of both the Test and Stimulus mice at the door as detected by the proximity sensor. For “Seeking” trials, social interactions lasting ≥4 seconds within the 12-second reward window were categorized as “Reciprocal”; otherwise, they were labeled “Nonreciprocal” (Movie 1–2). For “Exploring” trials, if interaction occurred for ≥3 seconds within the final 8 seconds of the reward period, the trial was classified as a “Late interaction”; otherwise, as “No interaction” (Figure 3A, Movie 3–5). For the seeking trials across a 9-day social stimulus period, 50.4% were nonreciprocal interactions where the Stimulus mouse did not approach, while 34.9% were reciprocal interactions (Figure 3B). This highlights a nuance of all social operant tasks: the Stimulus mouse is not obligated to reciprocate when the Test mouse initiates a social interaction. The proximity sensors allowed us to quantify this critical feature of the task.

**Figure 3.**
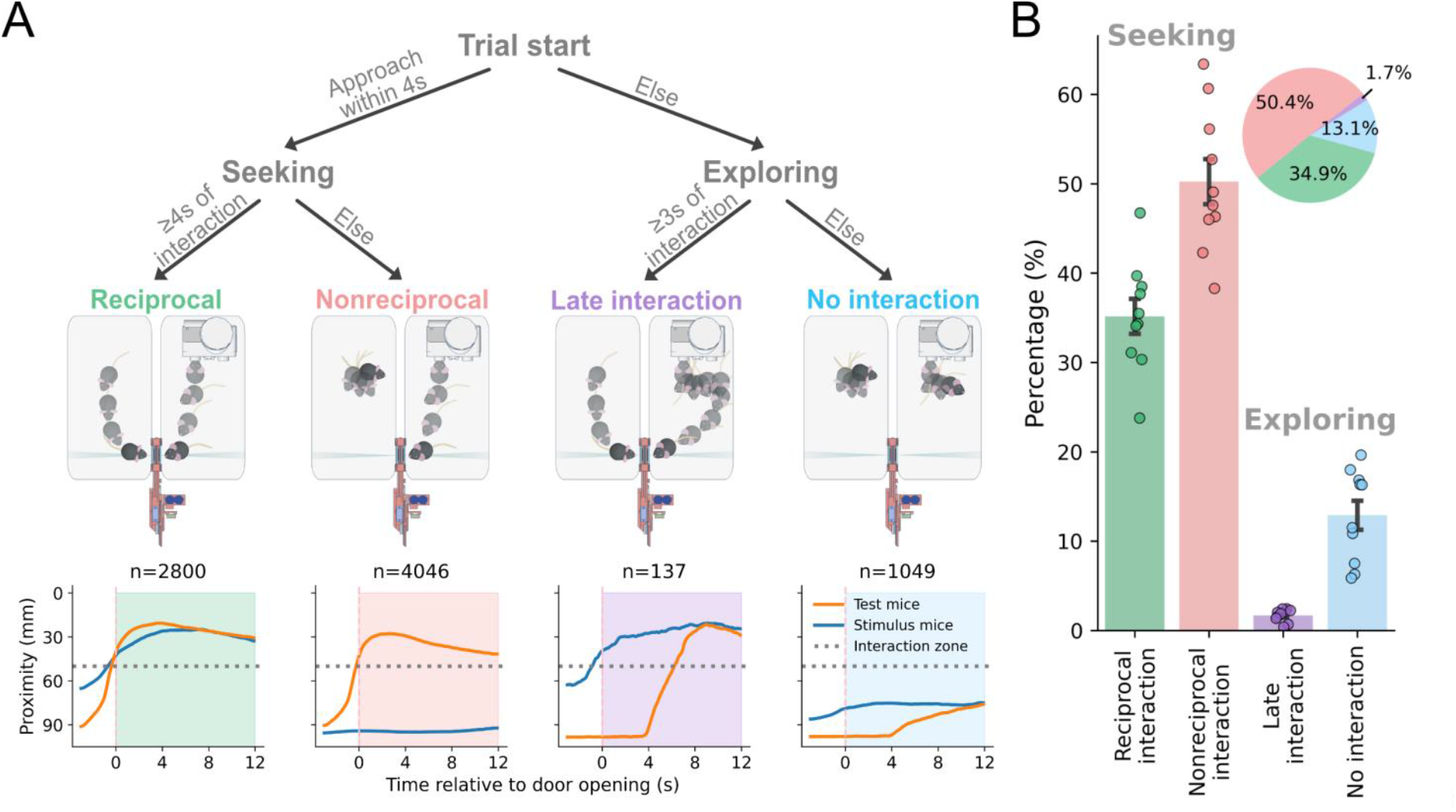
Post-door-opening social interactions can be categorized based on reciprocity and timing. (A) (top) Flowchart outlining the classification logic for social trials. Each trial begins when the door opens. If the test mouse approaches social door within 4 seconds, the trial is labeled as seeking; otherwise, it is classified as exploring. Seeking trials are further divided based on the duration and timing of interaction: reciprocated (⩾4 seconds of interaction within the first 12 seconds), not reciprocated, late interaction (<3 seconds of interaction during the last 8 seconds), or no interaction. (middle) Representative examples of mouse trajectories for each of the four outcome categories: reciprocal, nonreciprocal, late interaction, and no interaction. (bottom) Average proximity traces (± SEM) showing the distance between test and stimulus mice over time in each category. The area above the gray dashed line indicates the interaction zone. Reciprocal trials show sustained mutual proximity, while other categories reflect one-sided or delayed social responses. (B) Distribution of all classified trials (N = 10 mice). Most seeking events were nonreciprocal (50.4%) or reciprocal (34.9%), while exploring trials primarily resulted in no interaction (13.1%) or late interaction (1.7%). Pie chart shows overall proportions of each category across all trials.

### The circadian rhythm of social seeking is delayed relative to the feeding rhythm

We quantified the circadian rhythms of feeding and social seeking behaviors in the task simultaneously. The acrophase (timing of the peak) of the social seeking rhythm was delayed by 1.93 ± 0.25 hours relative to the feeding rhythm (N = 14, T = 0.0, p = 9.79 × 10^-4^, Wilcoxon signed-rank test) (Figure 4A–B), suggesting that mice prioritize caloric needs before initiating social engagement. Consistent with this idea, individual mice also shifted their feeding rhythms 1.16 ± 0.32 hours earlier when the target cage contained a Stimulus mouse, compared to when the same mouse was tested in the empty cage condition (N = 11, T = 5.0, p = 9.77 × 10^-3^) (Figure 4C–D). This demonstrates how the presence of a social partner can influence the temporal organization of other daily activities.

**Figure 4.**
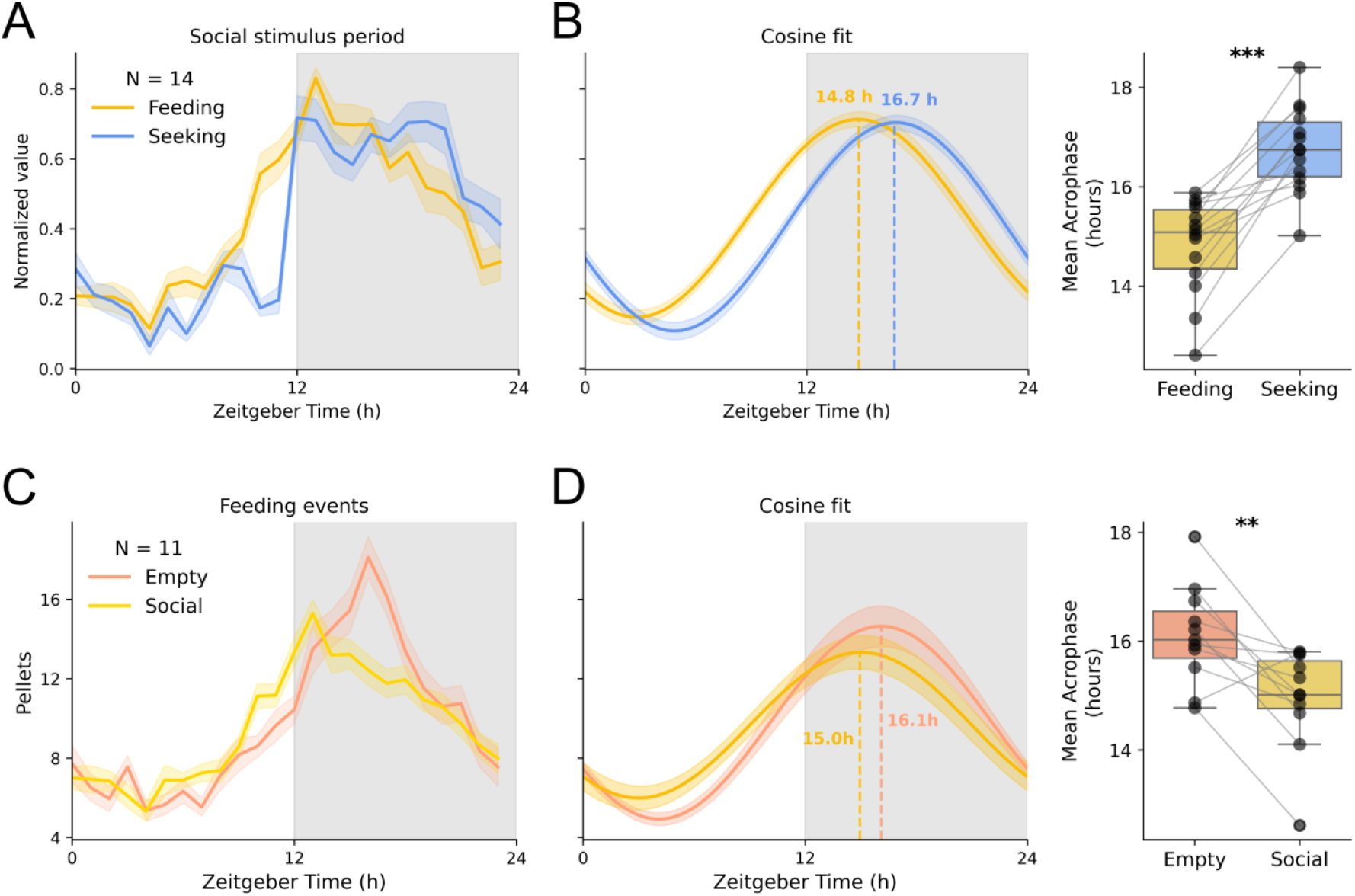
Social seeking peaks later than feeding, while social context shifts the timing of feeding behavior. (A) Daily patterns of feeding and social seeking during the social stimulus phase (N = 14). Both behaviors ramp up at dark onset (gray shading), but their peaks occur at different times. ZT0 corresponds to 6:00 AM (lights-on); ZT12 = 18:00PM (lights-off). (B) Cosine curve fits (left) and circular plot of individual acrophases (right) show that feeding peaks significantly earlier than social seeking (mean acrophase: 14.8 h vs. 16.7 h, p = 9.79 × 10^-4^, Wilcoxon signed-rank test). (C) Time course of feeding events across empty cage and social stimulus days (N = 11). ZT0 corresponds to 6:00 AM (lights-on); ZT12 = 18:00PM (lights-off). (D) Cosine fits and circular plots show a significant phase advance in feeding during social stimulus exposure compared to the empty cage condition (mean acrophase: 16.1 h vs. 15.0 h, p =9.77 × 10^-3^, Wilcoxon signed-rank test). Gray shading indicates the dark phase (ZT 12-24); shaded curves show mean ± SEM. Stats: Wilcoxon signed-rank test. Asterisks indicate statistical significance: p < 0.01 (**) and p < 0.001 (***). All tests were two-tailed, and multiple comparisons were corrected using the Bonferroni method where applicable.

### Dopaminergic impacts on social seeking

We validated the sensitivity of our behavioral assay to pharmacological manipulation of dopamine signaling, which has been widely implicated in motivated behavior and social reward processing^16–18^. Following the schematic in Figure 5A, we found that dopamine receptor antagonists significantly affected social and nonsocial behaviors (Figure 5B–E). Friedman tests revealed significant treatment effects on social seeking (N = 14, χ^2^(2) = 6.00, p = 0.0498), feeding (χ^2^(2) = 10.86, p = 4.39 × 10^−3^), and overall activity (χ^2^(2) = 9.85, p = 7.28 × 10^-3^), but not on exploring behavior (χ^2^(2) = 4.25, p = 0.12). Post-hoc tests revealed that SCH23390 selectively reduced social seeking (T = 13.0, p = 0.0213, Holm-Bonferroni-corrected), without affecting general locomotion (T = 39.0, p = 0.68) or feeding (T = 38.0, p = 0.39). In contrast, haloperidol significantly reduced social seeking (T = 6.5, p = 0.0213), overall activity (T = 0.0, p = 4.88 × 10^-4^), and feeding (T = 3.0, p = 1.22 × 10^-3^), suggesting a broad suppression of behavior rather than a specific social deficit.

**Figure 5.**
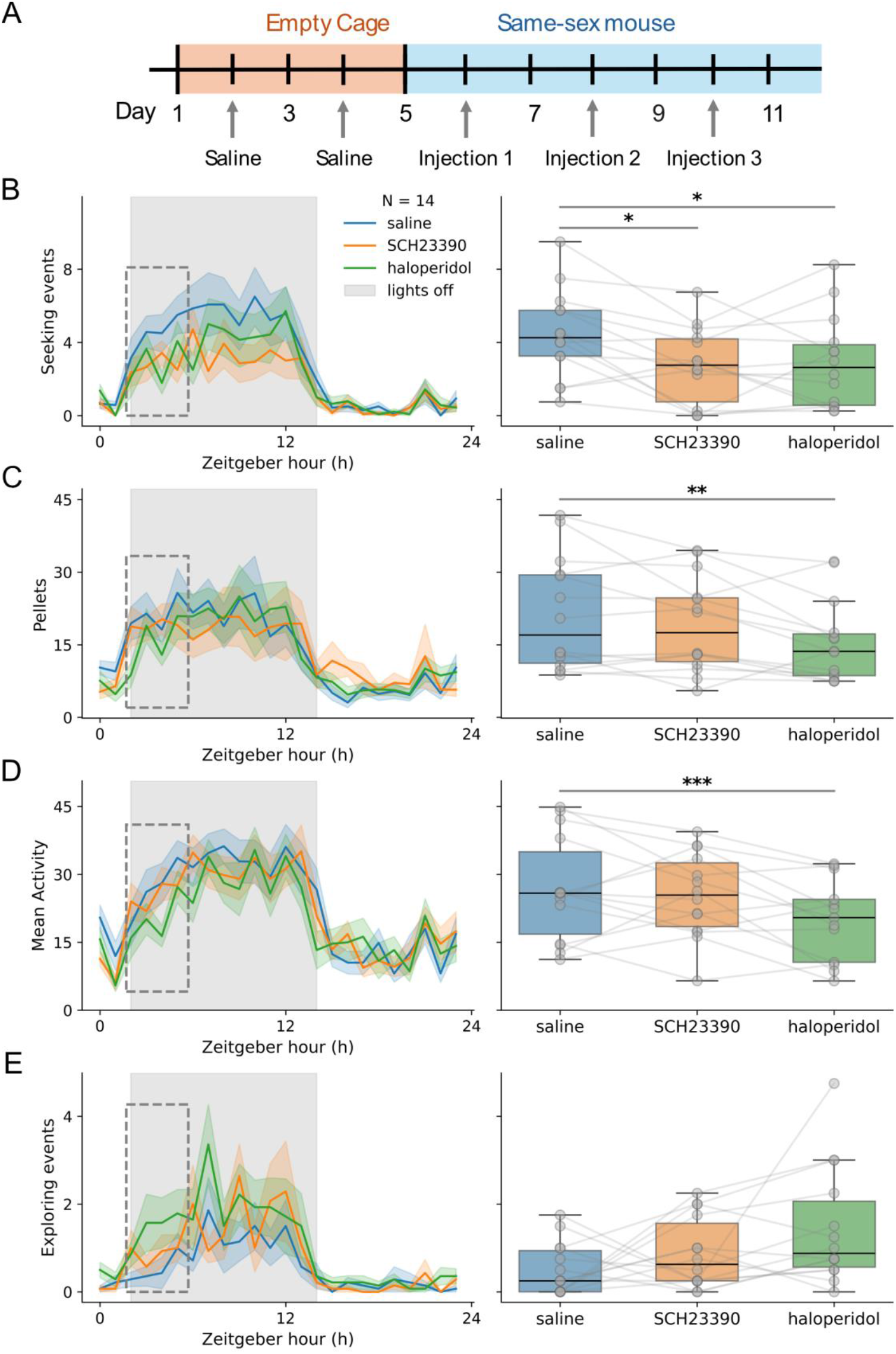
Social seeking behavior is selectively reduced by dopamine receptor antagonists. (A) Experimental timeline and counterbalanced drug design. Mice received three injections (saline, SCH23390, haloperidol) in randomized order during same-sex social stimulus exposure days. (B) Left: Time course of social seeking events over 24 h. ZT0 = 4:00 PM (lights on); ZT12 = 4:00 AM (lights off). Right: Quantification during the first 4 h of the dark phase (dashed box). SCH23390 (p = 0.021, Wilcoxon signed-rank test with Holm– Bonferroni correction) and haloperidol (p = 0.021) significantly reduced seeking compared to saline. (C) Feeding events across 24 h. (D) Total activity across 24 h. (E) Exploration events following door opening across drug conditions. Gray shading indicates the dark phase (ZT 12–24); shaded curves show mean ± SEM. Stats: Wilcoxon signed-rank test. Asterisks indicate statistical significance: p < 0.05 (*), p < 0.01 (**), and p < 0.001 (***). All tests were two-tailed, and multiple comparisons were corrected using the Bonferroni method where applicable.

## Discussion

This study introduces a novel home-cage social operant task that complements existing tests of social motivation. Our system consists of a battery-powered, sliding door positioned between two cages that can be opened by the Test mouse with a nose-poke. When the door opens, it allows limited social interaction (visual sensing and sniffing through a steel grate) between the Test mouse and the Stimulus mouse in the neighboring cage. By testing mice in their home environments, our system minimizes stress, reduces experimenter intervention, and enables continuous monitoring of social behavior over weeks.

Mice learned to nose-poke to open the door within one day, as evidenced by a significant increase in operant responding on the first day that a mouse was introduced to the neighboring cage, vs. when the neighboring cage was empty. This is consistent with prior findings showing that rodents are motivated to work for social access^8,9,11^. Video tracking provides a valuable approach for quantifying social behaviors in rodent operant tasks^11,19^. However, due to the length of our experiments (1-2 weeks), this analysis of social interactions would have been challenging with video-based analyses, highlighting the benefit of proximity sensors for continuous behavioral quantification over multiple days. While we used FED3 nosepoke ports in this experiment to record operant effort and trigger door opening, the system is not dependent on FED3. For researchers not focused on feeding behavior, the nosepoke module can be replaced with alternative input formats, such as capacitive touch sensors. This flexibility allows adaptation of the task to a broad range of experimental needs.

In typical social approach and operant paradigms, stimulus animals are often confined in small, restrictive chambers that may artificially constrain or force social proximity. In such contexts, test animals may initiate interaction not due to genuine social motivation, but in response to distress cues or escape attempts from the stimulus animal, raising the possibility that social contact initiated by the test animals reflects empathic or rescue-driven responses rather than voluntary social engagement. Indeed, mice have been shown to exhibit empathy-like behaviors toward distressed conspecifics^20^, and their behavior is sensitive to the movements of other animals^21^. Our paradigm differs by allowing the Stimulus animal free movement within an adjacent cage, enabling voluntary interaction and minimizing confounding stress responses. Using ToF proximity sensors, we classified social interactions and found that about half of the social seeking attempts by the Test mouse did not result in a reciprocated social interaction, as the Stimulus mouse never approached the door. These findings underscore a key but commonly overlooked issue of social interaction in rodents: the challenge of aligning motivational states between Test and Stimulus animals. This temporal desynchrony may arise from individual variability in internal state or competing behavioral priorities, and it should be carefully considered in the design and interpretation of social behavior experiments.

Our system also revealed that social seeking follows a circadian rhythm that peaks approximately two hours after the peak of the feeding rhythm. Interestingly, adding a social partner also shifted the timing of feeding behavior earlier by ∼1 hour, relative to the rhythms in the same mice without a social partner, suggesting that social drive can impact the feeding rhythm. One possible explanation for this finding is that the task and the associated social cues influenced the endocrine regulation of feeding, which in turn modulates the circadian clock governing metabolic rhythms^22^. Future experiments can incorporate hormonal assays to test this hypothesis and clarify the underlying mechanisms.

Several researchers have linked social motivation to the dopamine system. For instance, multiple researchers have found that the activity of dopamine neurons increases while animals are performing social behaviors^16–18,23,24^. In a critical finding, Solié et al. (2022) used a social operant task to determine that ventral tegmental area dopamine (VTA^DA^) neurons encode a “social prediction error” which can guide learning about social outcomes^24^, complementing a large literature showing “reward prediction errors” for many distinct outcomes in VTA^DA^ neurons ^25^. This work has also led to understanding the molecular underpinnings of dopamine function and how these mechanisms can be altered in ASD. For instance, mice that are deficient in SHANK3 have reduced VTA^DA^ neuron activity, which can explain their social deficits^11,17^. We were motivated by this prior literature to test if pharmacological manipulation of the dopamine system could alter operant behavior in the homecage social operant test. Consistent with the prior work on this topic, we found that D1 receptor blockade via systemic SCH23390 administration selectively impaired social seeking without affecting feeding or general activity levels. In contrast, D2 receptor blockade via systemic haloperidol administration suppressed social behavior, but also feeding and activity levels, consistent with its broader motor-suppressive effects. These findings are consistent with reports that SCH23390 selectively impairs social learning and social interaction without disrupting feeding^26,27^. Haloperidol, on the other hand, exerts more widespread effects on social behavior^28^, locomotion, and feeding^29^. A limitation of these experiments is that systemic drug administration lacks regional and receptor-type selectivity. While dopaminergic receptors in the nucleus accumbens are known to play key roles in social approach and motivational drive^23^, systemic delivery likely influences multiple brain regions and receptor populations beyond this target. Investigating circuit and cell-type-specific effects using this paradigm will be an important future direction.

Both sexes were used in all experiments, but identification of sex differences was not a goal of this study, and we are underpowered to properly assess them. Future work will be devoted to assessing sex differences in social motivation, including interaction types, social circadian rhythms, and competing drives for feeding and social seeking.

Together, these findings demonstrate that the home-cage social operant task is a powerful and versatile tool for investigating social motivation under naturalistic and low-stress conditions. Its compatibility with long-term, continuous monitoring makes it especially well-suited for studies involving neuromodulation, circadian dynamics, and individual variability in social behavior. In addition to replicating and extending key findings from traditional assays, this paradigm offers new insights into the temporal structure and neurochemical basis of social behavior. Moving forward, this system can be adapted to test developmental trajectories, sex differences, disease models, and circuit-specific manipulations, opening new avenues for research into the biological foundations of social interaction and their disruption in neuropsychiatric conditions.

## Acknowledgments

This work was supported by the National Institutes of Health: 1RM1MH138313-01 (AVK, SEM), DK136810 (AVK), DK138131 (AVK), and P50HD103525 to IDDRC at WashU. We thank Dr. Joseph Dougherty and Ms. Katherine McCullough for assistance in procuring mice, the Animal Behavior Subunit of the IDDRC at WashU for access to procedural space, Dr. Matthew Gaidica and the Department of Neuroscience NeuroTech Hub for assistance with apparatus construction, and members of the Kravitz and Creed labs for feedback on the manuscript.

## References

1. Insel, T. R. & Fernald, R. D. HOW THE BRAIN PROCESSES SOCIAL INFORMATION: Searching for the Social Brain*. Annual Review of Neuroscience 27, 697–722 (2004).

2. Cacioppo, J. T. & Hawkley, L. C. Perceived social isolation and cognition. Trends in Cognitive Sciences 13, 447–454 (2009).

3. Chevallier, C., Kohls, G., Troiani, V., Brodkin, E. S. & Schultz, R. T. The social motivation theory of autism. Trends in Cognitive Sciences 16, 231–239 (2012).

4. Insel, T. R. Rethinking schizophrenia. Nature 468, 187–193 (2010).

5. Fussner, L. M., Mancini, K. J. & Luebbe, A. M. Depression and Approach Motivation: Differential Relations to Monetary, Social, and Food Reward. J Psychopathol Behav Assess 40, 117–129 (2018).

6. Kupferberg, A., Bicks, L. & Hasler, G. Social functioning in major depressive disorder. Neuroscience & Biobehavioral Reviews 69, 313–332 (2016).

7. File, S. E. & Seth, P. A review of 25 years of the social interaction test. Eur J Pharmacol 463, 35–53 (2003).

8. Martin, L. & Iceberg, E. Quantifying Social Motivation in Mice Using Operant Conditioning. JoVE (Journal of Visualized Experiments) e53009 (2015) doi:10.3791/53009.

9. Hu, R. K. et al. An amygdala-to-hypothalamus circuit for social reward. Nat Neurosci 24, 831–842 (2021).

10. Ramsey, L. A., Holloman, F. M., Lee, S. S. & Venniro, M. An operant social self-administration and choice model in mice. Nat Protoc 18, 1669–1686 (2023).

11. Maloney, S. E. et al. A comprehensive assay of social motivation reveals sex-specific roles of autism-associated genes and oxytocin. Cell Reports Methods 3, (2023).

12. Bailey, J. Does the Stress of Laboratory Life and Experimentation on Animals Adversely Affect Research Data? A Critical Review. Altern Lab Anim 46, 291–305 (2018).

13. Sorge, R. E. et al. Olfactory exposure to males, including men, causes stress and related analgesia in rodents. Nat Methods 11, 629–632 (2014).

14. Matikainen-Ankney, B. A. et al. An open-source device for measuring food intake and operant behavior in rodent home-cages. eLife 10, e66173 (2021).

15. Refinetti, R., Lissen, G. C. & Halberg, F. Procedures for numerical analysis of circadian rhythms. Biol Rhythm Res 38, 275–325 (2007).

16. Gunaydin, L. A. et al. Natural Neural Projection Dynamics Underlying Social Behavior. Cell 157, 1535–1551 (2014).

17. Bariselli, S. et al. SHANK3 controls maturation of social reward circuits in the VTA. Nat Neurosci 19, 926–934 (2016).

18. Manduca, A. et al. Dopaminergic Neurotransmission in the Nucleus Accumbens Modulates Social Play Behavior in Rats. Neuropsychopharmacol 41, 2215–2223 (2016).

19. Raymond, J. S., Rehn, S., James, M. H., Everett, N. A. & Bowen, M. T. Sex differences in the social motivation of rats: Insights from social operant conditioning, behavioural economics, and video tracking. Biol Sex Differ 15, 57 (2024).

20. Ueno, H. et al. Empathic behavior according to the state of others in mice. Brain and Behavior 8, e00986 (2018).

21. Ueno, H. et al. Conformity-like behaviour in mice observing the freezing of other mice: a model of empathy. BMC Neurosci 21, 19 (2020).

22. Tsang, A. H., Astiz, M., Friedrichs, M. & Oster, H. Endocrine regulation of circadian physiology. (2016) doi:10.1530/JOE-16-0051.

23. Dai, B. et al. Responses and functions of dopamine in nucleus accumbens core during social behaviors. Cell Reports 40, (2022).

24. Solié, C., Girard, B., Righetti, B., Tapparel, M. & Bellone, C. VTA dopamine neuron activity encodes social interaction and promotes reinforcement learning through social prediction error. Nat Neurosci 25, 86–97 (2022).

25. Schultz, W. Reward prediction error. Curr Biol 27, R369–R371 (2017).

26. Matta, R., Tiessen, A. N. & Choleris, E. The Role of Dorsal Hippocampal Dopamine D1-Type Receptors in Social Learning, Social Interactions, and Food Intake in Male and Female Mice. Neuropsychopharmacol 42, 2344–2353 (2017).

27. Choleris, E., Clipperton-Allen, A. E., Gray, D. G., Diaz-Gonzalez, S. & Welsman, R. G. Differential Effects of Dopamine Receptor D1-Type and D2-Type Antagonists and Phase of the Estrous Cycle on Social Learning of Food Preferences, Feeding, and Social Interactions in Mice. Neuropsychopharmacol 36, 1689–1702 (2011).

28. Redolat, R., MIÑARRO, J., Simon, V. M. & Brain, P. F. Influences of Haloperidol and Sulpiride on Social Behavior of Female Mice in Interactions with Anosmic Males. 79–85 (1991).

29. Huang, A. C. W., Shyu, B.-C. & Hsiao, S. Dose-dependent dissociable effects of haloperidol on locomotion, appetitive responses, and consummatory behavior in water-deprived rats. Pharmacology Biochemistry and Behavior 95, 285–291 (2010).

