## Supplemental Figure for "A novel homecage operant paradigm reveals circadian and behavioral dynamics of social motivation in mice"

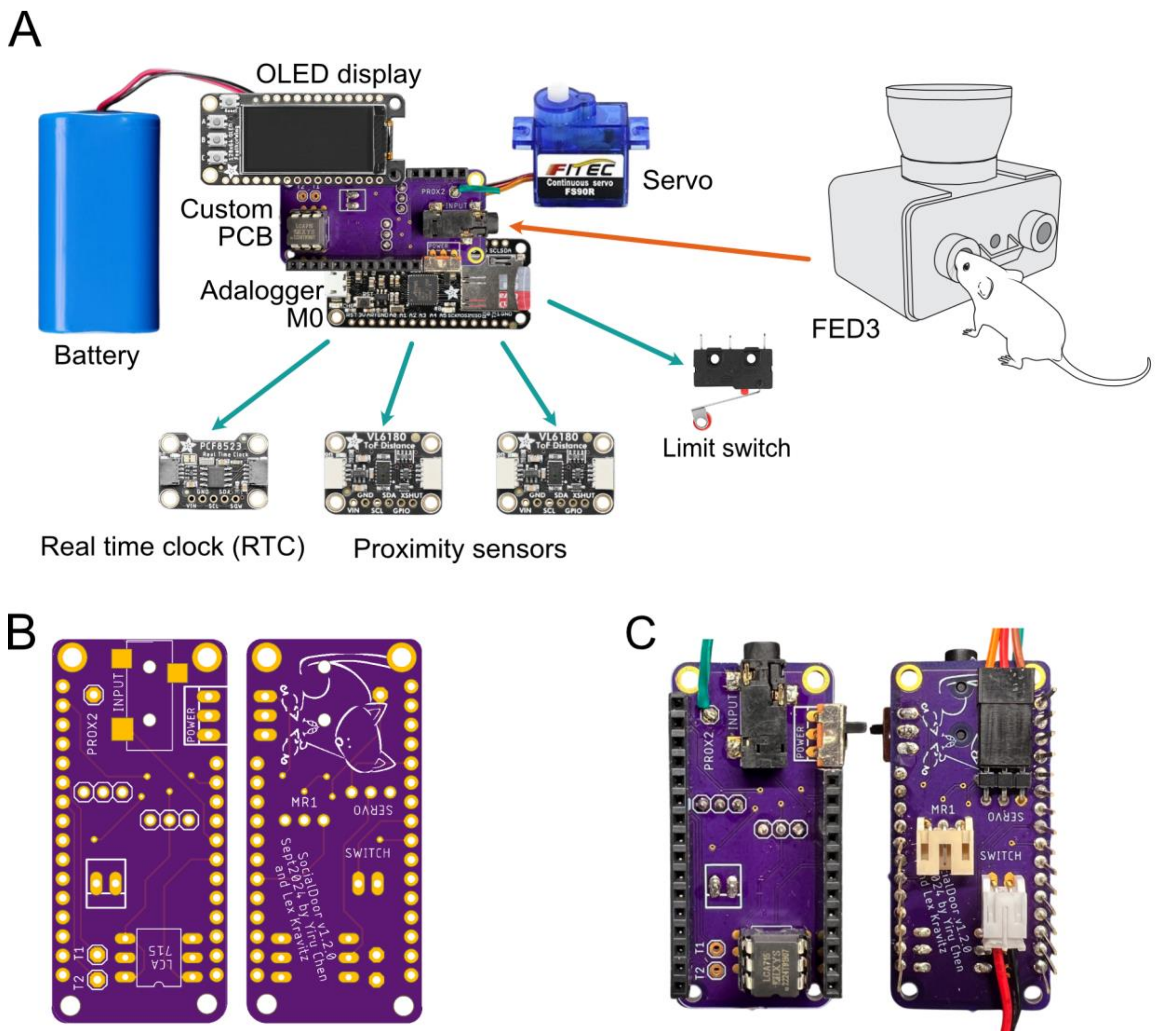


**Supplemental Figure 1. Circuit design and hardware components used in the homecage social operant system.**

(A) Overview of the complete electronic system used to control the homecage social task (see also Figure 1A–B). The core unit is built around an Adalogger M0 microcontroller mounted on a custom printed circuit board (PCB), which interfaces with an OLED display, battery, and external sensors. These include a real-time clock (RTC) module for timekeeping, two VL6180X time-of-flight proximity sensors to detect the positions of the test and stimulus mice, respectively, and a limit switch for door position feedback. A servo motor controls the movement of the sliding door in response to behavioral triggers. The system operates independently and can interface with a FED3 device for nosepoke input.

(B) Front and back views of the custom PCB prior to assembly.

(C) Fully assembled PCB with components and connectors soldered in place. Connectors support plug-and-play attachment of the servo motor, limit switch, and other external modules.

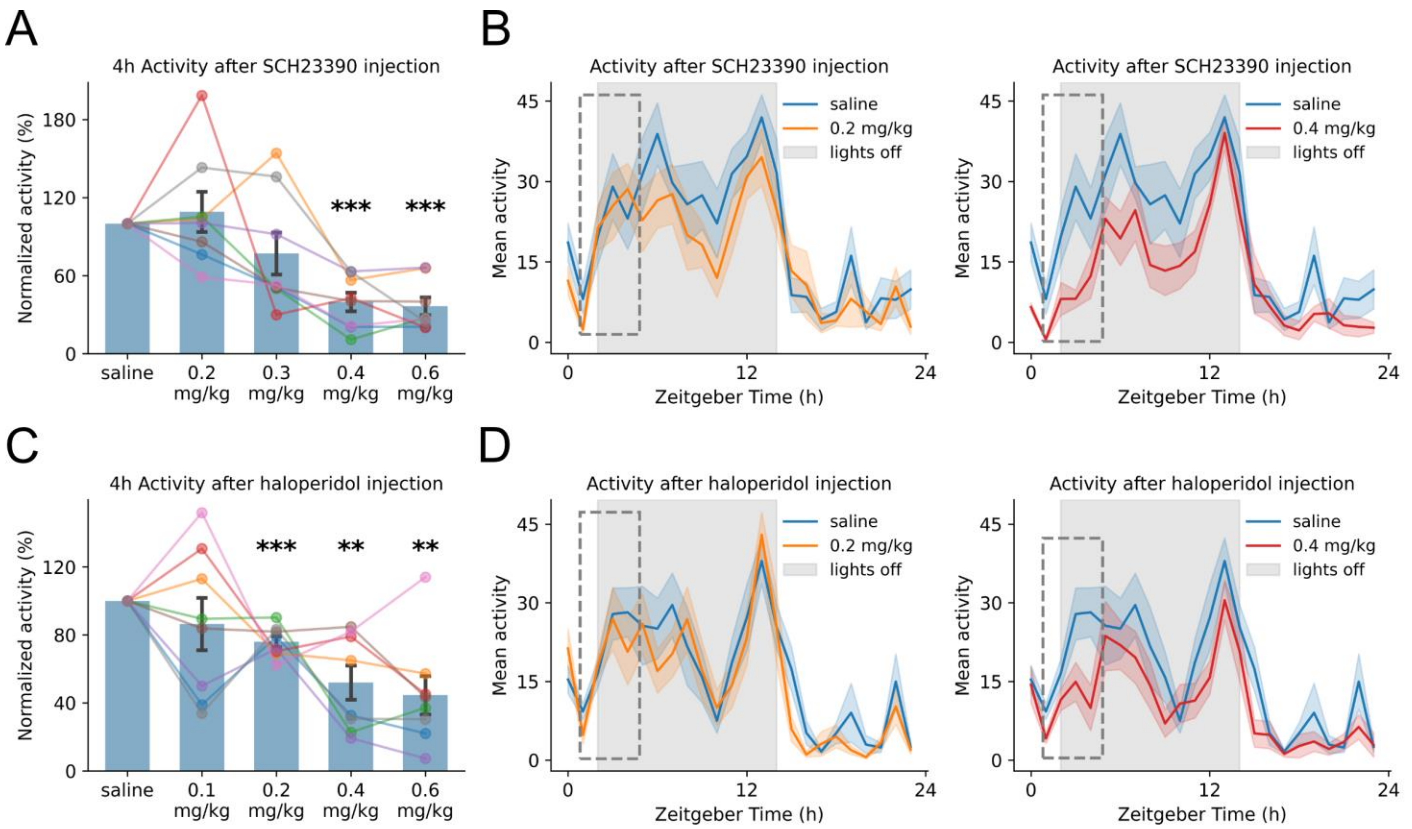


**Supplemental Figure 2. Dose-dependent effects of SCH23390 and haloperidol on locomotor activity.**

(A) Locomotor activity during the 4-hour period following SCH23390 injection, expressed as a percentage of each mouse's saline baseline. While lower doses had no significant effect, activity was significantly reduced at 0.4 mg/kg (N = 8, $p = 2.11 \times 10^{-4}$) and 0.6 mg/kg ($p = 1.35 \times 10^{-4}$), as determined by paired t-tests with Holm-Bonferroni correction.

(B) 24-hour activity profile following SCH23390 (0.2 mg/kg or 0.4 mg/kg) or saline injection. The gray dashed box corresponds to the 4-hour window shown in panel A. Locomotor activity was reduced during the early post-injection period at 0.4 mg/kg, but not 0.2 mg/kg. Overall circadian rhythmicity was preserved at both doses.

(C) Activity following haloperidol injection, normalized to saline. Activity declined in a dose-dependent manner, with significant reductions at 0.2 mg/kg (N = 8, $p = 5.80 \times 10^{-4}$), 0.4 mg/kg (p = 0.0052), and 0.6 mg/kg (p = 0.0052), as determined by paired t-tests with Holm-Bonferroni correction.

(D) 24-hour activity profile following haloperidol (0.2 mg/kg or 0.4 mg/kg) or saline injection. The gray dashed box corresponds to the 4-hour window shown in panel A. Both doses suppressed early activity, while diurnal rhythmicity remained intact.

Shaded regions represent SEM. Gray backgrounds indicate the dark phase (ZT2-14). ZT0 corresponds to 4:00 PM (lights-on); ZT12 = 4:00 AM (lights-off).

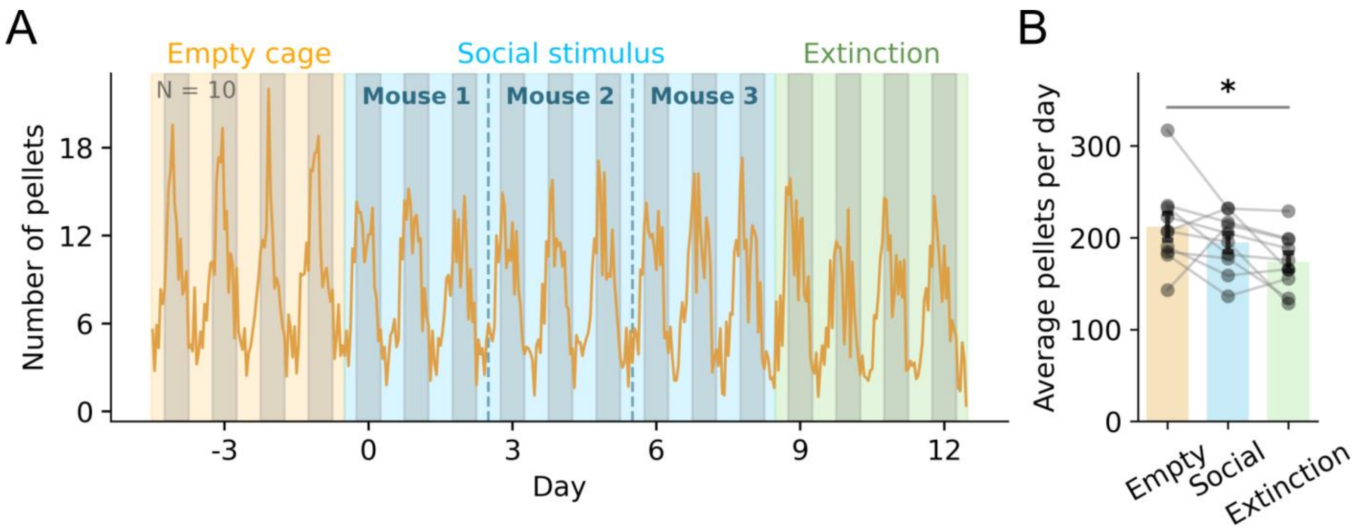


**Supplemental Figure 3. Feeding behavior is stable during social exposure but decreases during extinction.**

(A) Hourly pellet retrieval across the 17-day operant experiment. Each phase is represented by a different color: Empty cage (orange), Social stimulus (blue), and Extinction (green). Gray shading indicates the dark phase. The social period involved sequential exposure to three novel same-sex conspecifics (dashed lines).

(B) Average number of pellets retrieved per day during each phase. Gray lines connect data from individual animals. Pellet retrieval significantly decreased during extinction compared to the empty cage period (N = 10, $p = 0.038$, post-hoc test with Holm-Bonferroni correction); no other comparisons were significant.

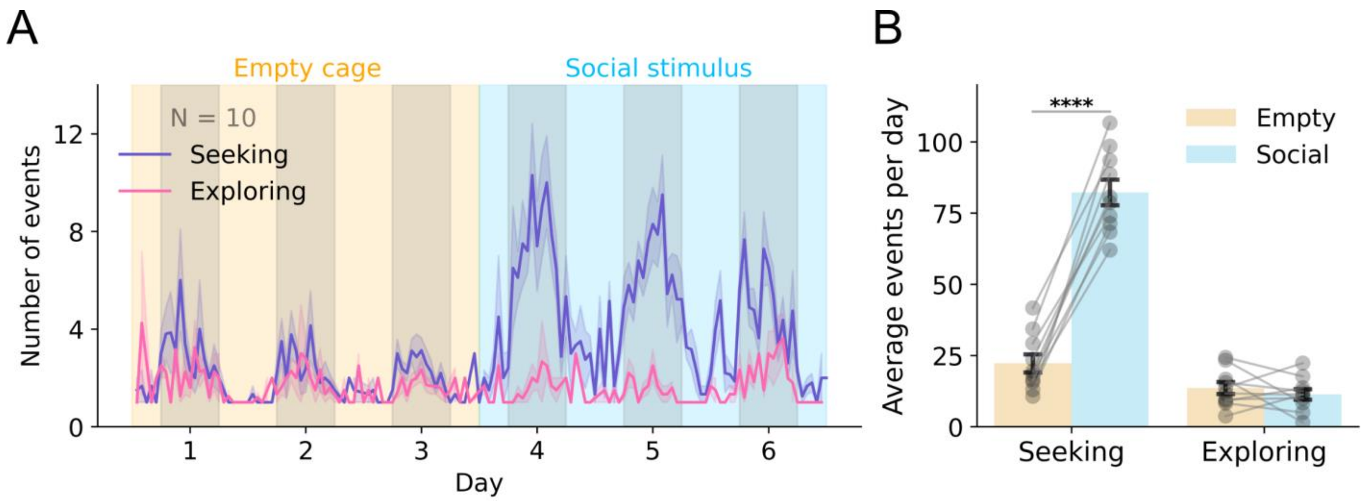


**Supplemental Figure 4. Social context selectively enhances seeking behavior and promotes operant learning.**

(A) Same dataset as Figure 1E, now separated into seeking (blue) and exploring (pink) events based on behavioral classification following door opening. Mice (N = 10) experienced alternating exposure to an empty cage (orange shading) or a same-sex social stimulus (blue shading) over 6 days. Seeking behavior increased prominently during social stimulus exposure, while exploring remained consistently low.

(B) Quantification of average daily events shows a significant increase in seeking during social stimulus days compared to empty cage ($p = 2.51 \times 10^{-7}$, paired t-test). Exploring behavior did not significantly differ between conditions ($p = 0.42$).

Asterisks indicate statistical significance: $p < 0.0001$ (****). Shaded regions indicate dark phase (lights off), and shaded lines represent mean ± SEM. Each dot represents one animal.
